# Fitness incentives to male fighters undermine fighting performance in intergroup contests

**DOI:** 10.1101/2024.05.09.593361

**Authors:** P.A. Green, D.W.E. Sankey, T. Collins, F. Mwanguhya, H. J. Nichols, M.A. Cant, F.J. Thompson

**Affiliations:** Department of Ecology, Evolution, and Marine Biology, University of California, Santa Barbara, CA 93106; Centre for Ecology and Conservation, College of Life and Environmental Sciences, University of Exeter, Penryn, Cornwall TR10 9FE, United Kingdom; Department of Ecology, Evolution, and Organismal Biology, Brown University, Providence, RI 02912; Department of Computer Science, University of Exeter, Exeter EX4 4QF, United Kingdom; Banded Mongoose Research Project, Queen Elizabeth National Park, Uganda; Department of Biosciences, Swansea University, Singleton Park Campus, Swansea, SA2 8PP, United Kingdom

**Keywords:** Intergroup conflict, social evolution, parochial altruism, animal contest, collective action

## Abstract

In humans and other animal societies, groups engage in intergroup conflicts over resources. The success of groups in these conflicts depends on individual contributions to collective fighting, yet individuals may have personal fitness incentives to defect rather than fight, which could undermine group performance. Here we test the hypothesis that personal fitness incentives affect intergroup conflict success in wild banded mongooses (*Mungos mungo*). In this species, intergroup fights are sometimes initiated by estrous females, who gain outgroup matings while their male group-mates contribute most of the fighting effort. We found that group fighting success was highest when a group’s females were in estrus, suggesting that, although females may initiate fights, their male group-mates seem motivated to chase away rival groups to defend their paternity. Surprisingly, we found that groups that won fights conceded more paternity to their rivals than groups that lost. In other words, behavioral “wins” did not always result in fitness “wins”. Younger males were more successful at attaining paternity between groups compared to within their own groups, suggesting that they may forego intergroup fighting to focus on intergroup mating. Overall, our results suggest that personal fitness incentives—here, in the form of paternity—vary widely among group members and can undermine rather than promote collective fighting performance. Such conflicts of interest are likely inherent in group combat and can contribute to variation in the frequency and costliness of intergroup violence.

## Introduction

Conflicts between groups, or “intergroup contests”, occur across diverse social-living taxa, from snapping shrimp (1) to birds (2, 3). Despite the fact that members of social-living groups may have disparate interests and motivation, group members are hypothesized to contribute to the collective good of defeating a rival group because winning comes with fitness rewards for participants. In formal models of intergroup conflict, for example, members from winning groups are assumed to gain direct reproductive benefits (4, 5). Observational studies of human societies find support for this assumption: for instance, male warriors from winning groups in societies from East Africa and South America gain fitness benefits in the form of marriages and resulting children (6, 7). In contrast to the evidence from human societies, there is very little understanding of the direct fitness consequences (i.e., parentage over offspring) of conflict outcomes in non-human animals. Instead, most studies quantify changes in related metrics like territory size (3, 8, 9), or show how contest frequency, not outcomes, affects reproduction or offspring survival (10–12). Quantifying the fitness consequences of intergroup conflict outcomes in non-human taxa is essential to understanding what motivates group members to contribute to a collective good, informing theories of the evolution of intergroup warfare. Here, we examine how fitness costs and benefits, in the form of paternity over offspring, relate to conflict outcomes in a model wild social mammal species, the banded mongoose (*Mungos mungo*).

## Results

Banded mongooses engage in frequent (13) and violent intergroup contests: they are one of a handful of social mammals (including lions, wolves, chimpanzees, and humans) in which intergroup conflict accounts for more than 10% of adult deaths of known cause (14). Our study population in Queen Elizabeth National Park, Uganda is composed of 10-12 social groups, each containing approximately 20 (median = 18, IQR = 9.25 (15)) adults as well as juveniles (offspring less than 6 months old (16)). Group sex ratios are male-biased, with an average of 1.8 males per female (15). All females within a group enter estrus within one week of each other, an event termed “group estrus”. Estrous females are closely mate-guarded by dominant males who are usually the oldest males in the group (15) and who obtain most within-group paternity, while younger males are mostly excluded from reproduction. On average, the three oldest males in groups sire 85% of juveniles per group breeding attempt (17).

At all times during the reproductive cycle, but especially during group estrus, groups engage in intergroup contests (10) (Fig. 1). Many contests begin because estrous females lead their group into a rival group’s territory (14, 18), triggering an intergroup contest during which the females can escape their within-group mate guard and mate with a male from the rival group. Females are thought to initiate intergroup contests and mate with rival group males because rival group males are less closely related to them than are their within-group mate guards (19). Juveniles conceived as a result of outgroup matings are more genetically heterozygous, heavier, and more likely to survive to adulthood (one year) than within-group offspring (20). While females appear to initiate contests to gain fitter offspring, males are the main participants, showing higher levels of aggression (18), paying more of the costs of injury and death (14), and being more important to contest success (13) as compared to females.

**Figure 1.**
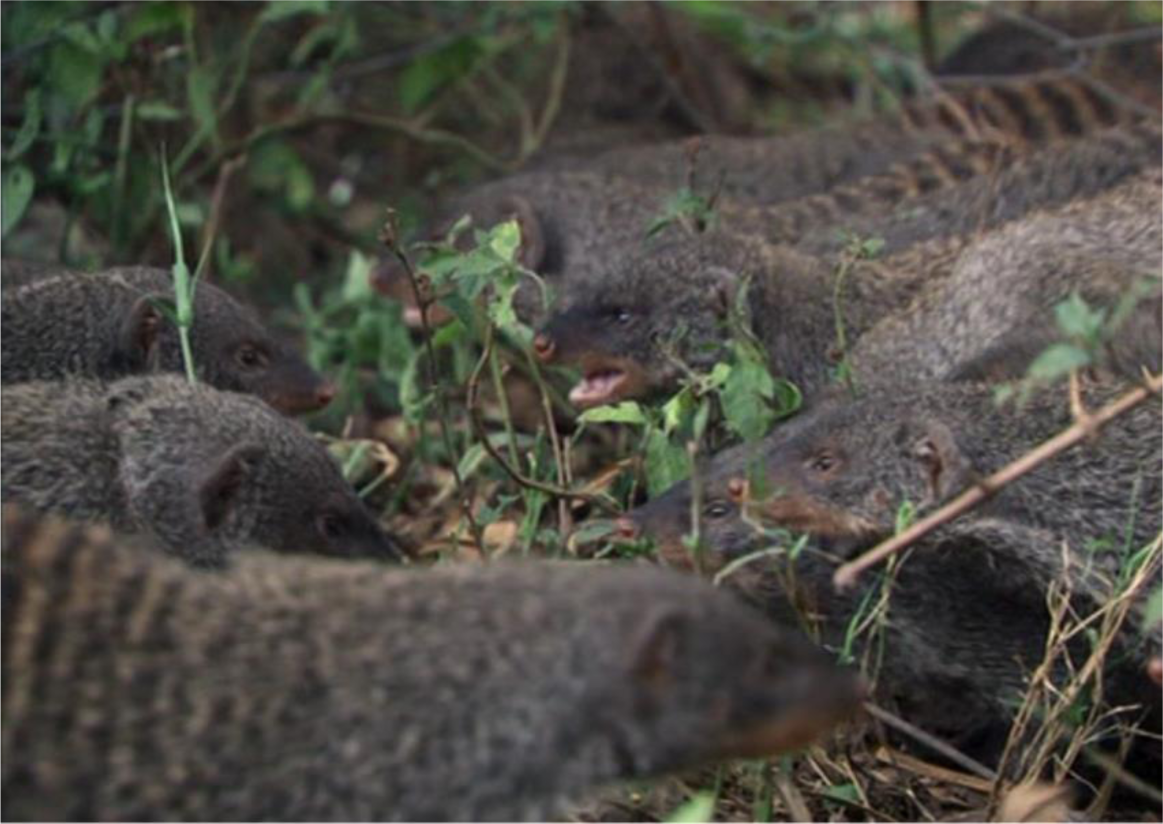
Banded mongooses during an intergroup contest. Although all group members can participate, males contribute more and pay more of the costs of competition. Photo: Mark MacEwen.

Our current understanding of this system, therefore, presents a conundrum: why do males participate in fights that females start but contribute little to? We tested the hypothesis that males are motivated to fight to defend their paternity; that is, males in focal groups are motivated to quickly dispel rival males from the site of the contest, thereby reducing the likelihood that these rival males mate with focal group estrous females and usurp focal male paternity.

We tested this hypothesis by adapting the framework of animal contest theory from its roots in dyadic (one-on-one) contests (21) to the case of intergroup fights (22). In the contest literature, contest success is defined as the ability of an individual or, in this case, of a group, to dispel its rival from the site of the contest (23). This success depends on two main factors: 1) resource holding potential (RHP), which is a measure of intrinsic fighting ability, and 2) the motivation to compete, based on the valuation of the contested resource, termed “resource value” or RV (21, 24–26). In banded mongoose societies, prior work has shown that group RHP is related to the number of males and age of the oldest male, which are the primary determinants of group fighting ability and predict a group’s success in dispelling its rival from the site of the contest (13). Our hypothesis predicts that group estrous status is a primary determinant of RV to male fighters. That is, if males from groups in estrus are motivated to displace rival males and maintain paternity over own-group offspring, then groups in estrus should be likely to win against groups not in estrus.

To test this prediction, we gathered data on the outcomes of 261 observed contests between banded mongoose groups. We also gathered data on each competing group’s estrous status at the time of the contest. We constructed a statistical model predicting the likelihood of contest success from three categories of relative group estrous status: neither competing group was in estrus, one group was and one group was not, or both groups were in estrus. When one group was in estrus and the other was not, the estrous group won 62% (64/104) of contests, while in other relative estrous scenarios, contest success averaged 49% (77/157 contests; MCMCglmm estimate mean of estrous effect on outcomes = 0.69, 95% CI = 0.08, 1.13, pMCMC = 0.03; Fig. 2; Table S1). In follow-up analyses, we found that the effect of group estrous on outcomes was stronger than that of a previously-established metric of group RHP, the relative age of the oldest male in the group, and that other potential metrics of resource value (number of females, number of juveniles, contest location) had little impact on contest outcomes (Fig. S1, Table S2, Table S3).

**Figure 2.**
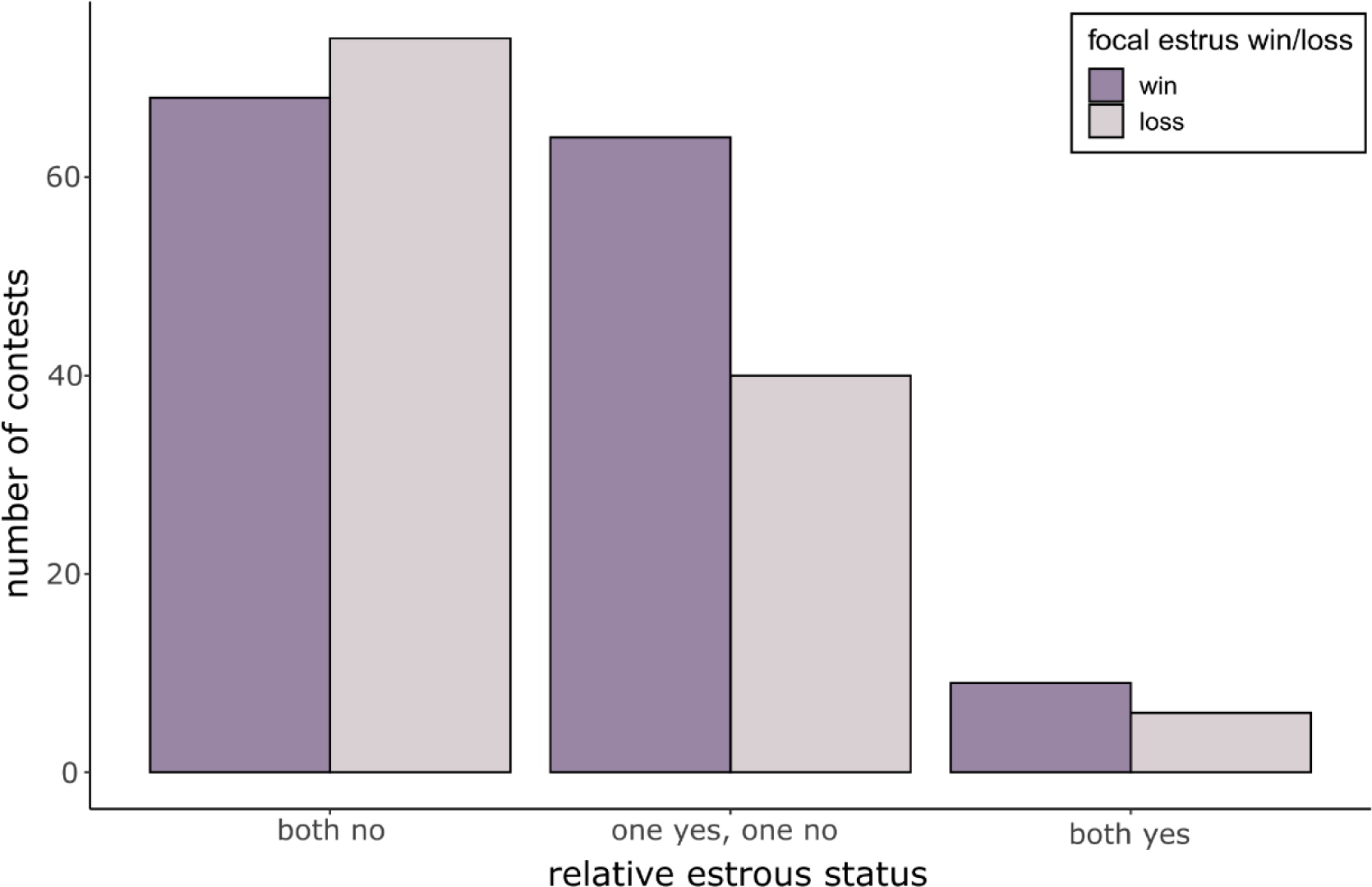
Female estrus status predicted contest outcomes. Barplot showing the number of contests won (dark purple bars) or lost (light purple bars) by groups according to their group estrous status and that of the rival group. Groups in group estrus fighting against groups not in group estrus (“one yes, one no” category) won 62% (64/104) of contests, a significantly higher winning rate than groups in other scenarios (Table S1).

If male fighting effort was motivated by the goal of defending paternity, our hypothesis further predicts that success in displacing rival groups should result in a maintenance of or (in scenarios where rival group females were also in estrus) gain of paternity over juveniles for in-group males. That is, winning in a behavioral sense should also result in winning in a fitness sense. This is assumed in theories of the evolution of warfare (4, 5) and is supported by observational studies of human societies, where winning intergroup fights comes with fitness benefits to participants (6, 7). To test this prediction in banded mongooses, we identified juveniles born after an intergroup contest and conceived during a time in which we recorded only one contest for the juveniles’ group in its estrous period (N = 197 juveniles in 48 contests). As such, we could be as confident as possible that, if these juveniles were sired by rival group males, this outgroup mating would likely have occurred during the contest in question. In total, 18% (36/197) of juveniles born after these contests were sired by males from rival groups. This outgroup siring was uneven across contests: in 25% of contests (12/48), more than half of the juveniles born in a given litter were sired by males from rival groups (mean ± sd = 61 ± 30%; range = 17% – 100% of juveniles born per litter). In the remaining 75% of contests, all juveniles were sired by ingroup males.

Counter to our prediction that winning groups retained or gained paternity, focal groups (randomly-assigned, see Methods) that won contests were more likely to lose paternity of their juveniles to males from the rival group, as compared to groups that lost contests. Focal groups that won contests lost paternity of 18 juveniles and gained paternity of 6, a net loss of paternity of 12 juveniles. By contrast, focal groups that lost contests lost paternity of only 6 juveniles and gained paternity of 6, a net of 0 (Fig. 3, Table S4). Winning and losing groups retained paternity of a similar number of juveniles, and results were broadly similar irrespective of relative estrous status (Table S3). A statistical model testing the effects of contest outcome (win/loss), relative estrous status, and their interaction on the gain or loss of paternity by focal groups showed that males from winning groups tended to lose paternity, compared to males from losing groups (β = – 1.23, SE = 0.80, χ^2^ = 2.36, P = 0.12; see Methods, Table S5). However, no effects in the model were statistically significant when controlling for litter, group, and contest identity. The same result held when analyzing the consequences of winning and losing on paternity of the proportion—instead of the absolute number—of juveniles sired in each litter; that is, while controlling for litter size (See Supplementary Information and Table S6). Overall, these results offer evidence that runs counter to assumptions of common theories of intergroup conflict (4, 5) by showing that winning an intergroup contest does not always result in a fitness benefit to males from winning groups.

**Figure. 3.**
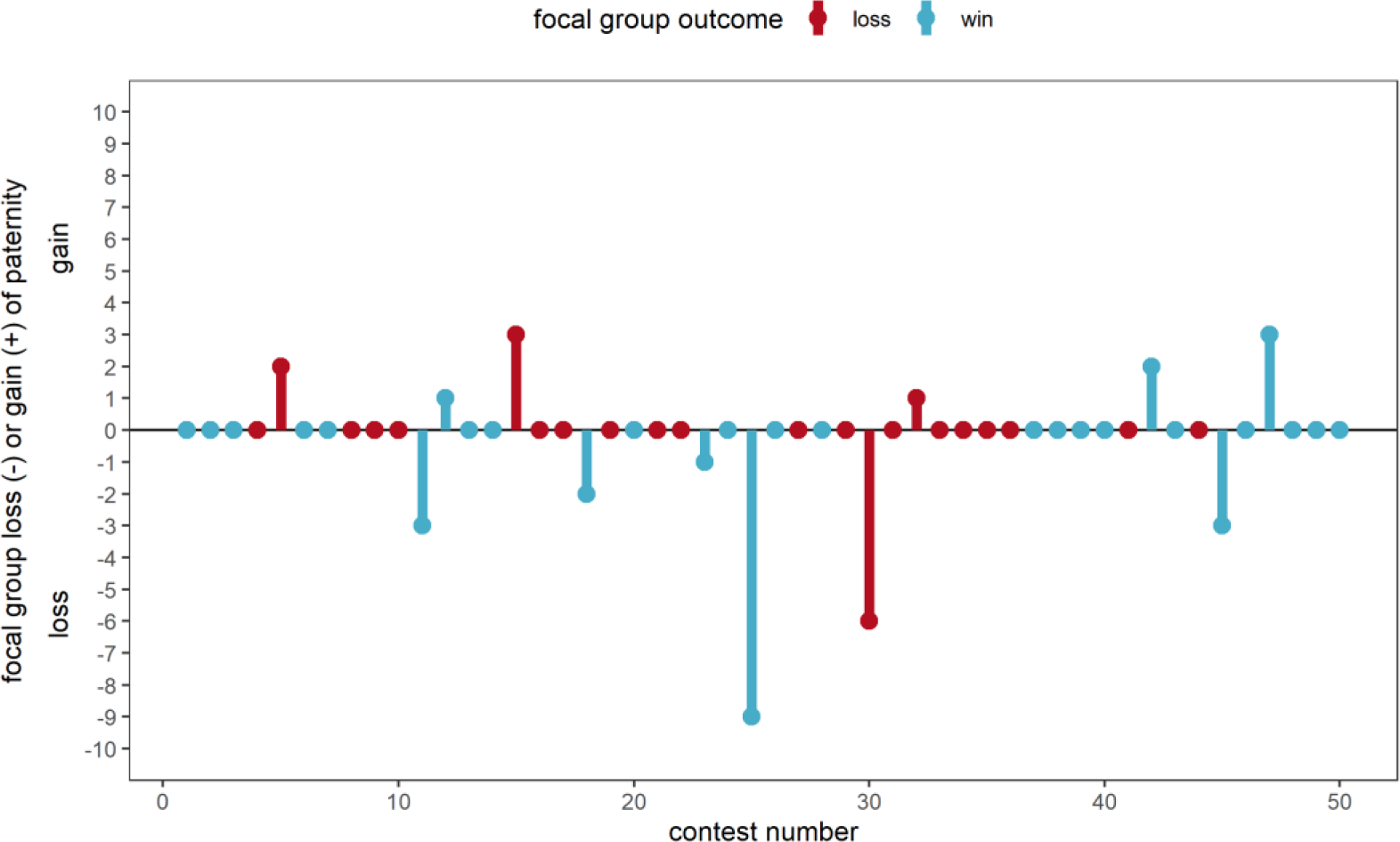
Males from winning groups experienced a net loss of paternity to males from losing groups. Points with connecting bars show, for each contest (x-axis), the number of juveniles of which paternity was gained (positive y-axis values) or lost (negative y-axis values) by focal groups (focal group identity was randomly assigned, see Methods; values of zero reflect retention of paternity by the focal group). Blue and red points and lines represent contests that the focal group won or lost, respectively. X-axis is ordered based on ranked date of contest (lower contest numbers occurred earlier in dataset). In 11 contests, litters born after contests contained juveniles with both focal and rival group sires; each of these contests has two contest numbers for clarity. Summary statistics are detailed in Table S4.

To further explore the fitness implications of intergroup contests to male group members, we tested which males in groups had the most to gain by mating rather than fighting. That is, which members might be least likely to contribute to the collective good of winning the fight. Within-group paternity in banded mongooses is usually monopolized by the oldest males in each group: a prior analysis focusing on within-group siring found that the oldest males in groups sire the greatest proportion of juveniles per litter (17). We built a statistical model replicating this prior analysis while adding data on juveniles sired by outgroup males, i.e., those sired during intergroup conflicts. Contrasting the within-group fitness advantage of older males, our results show that younger and older males sired an equal proportion of juveniles per litter during intergroup conflicts (GLMM estimate for interaction term of effect of sire age rank and paternity category = 0.04, SE = 0.02, χ^2^ = 4.90, df = 1, P = 0.03; Fig. 4, Table S7). Therefore, younger males in groups seem able to gain paternity during intergroup conflict that they could not gain through within-group mating.

**Figure 4.**
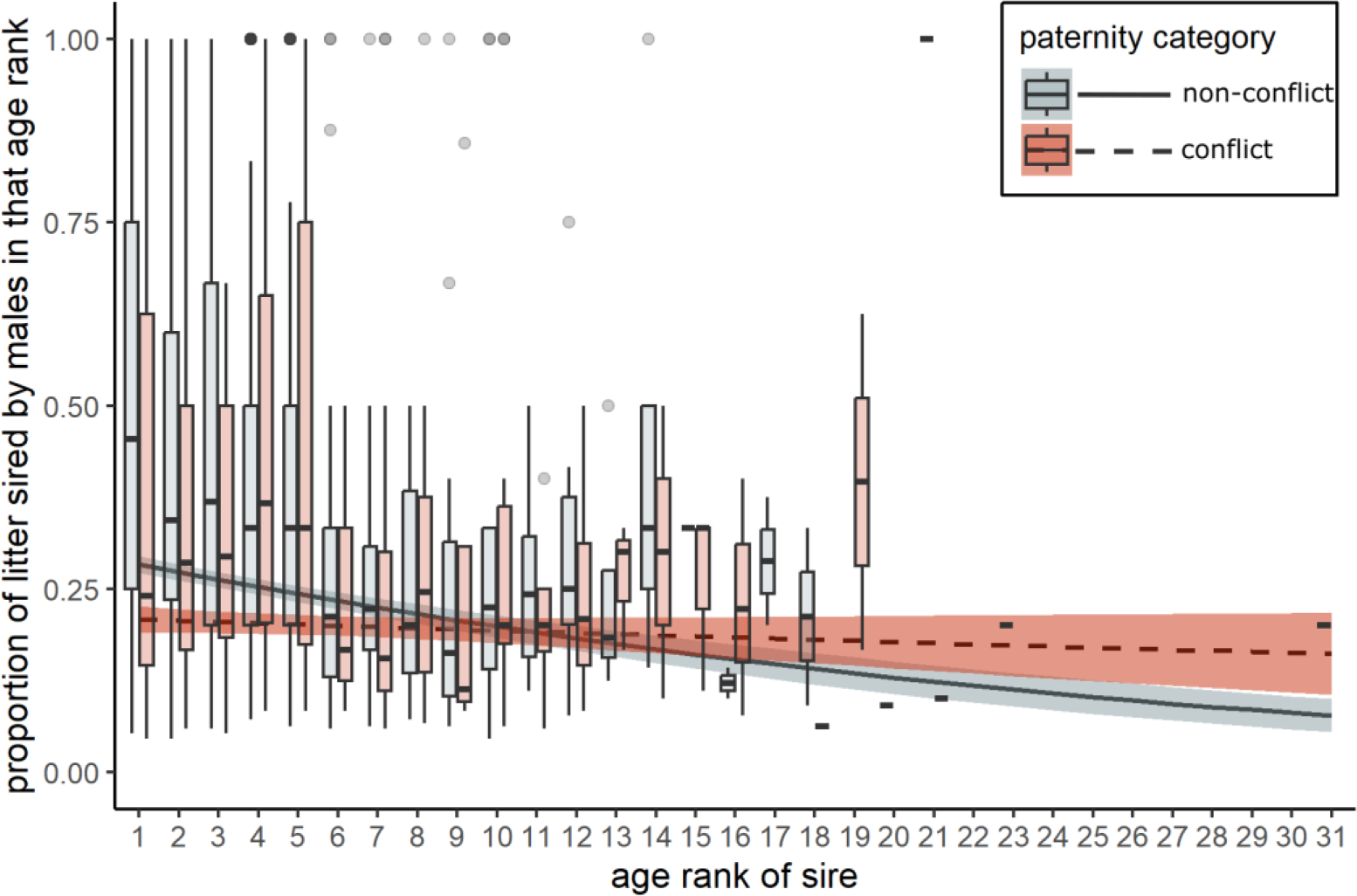
Paternity outside of conflict was monopolized by older males in groups, but all males had similar paternity over juveniles sired during intergroup conflict. Boxplot showing the proportion of juveniles sired by males in each age rank (1 = oldest male in their own group). Gray boxes show paternity during non-conflict scenarios; orange boxes show paternity during intergroup conflict. Lines with error ribbons show predicted values from the interaction term of a binomial GLMM predicting the proportion of litters sired by age rank, non-conflict (solid line with gray ribbon) or conflict (dashed line with orange ribbon) paternity, and their interaction.

## Discussion

Previous studies of intergroup conflict in non-human animals have measured how contest outcomes are related to fitness proxies such as territory residency (2, 3, 8, 27) and movement patterns (9, 28), or have focused on how conflict frequency influences birth rates and offspring survival (10–12). Studies of the direct fitness consequences (e.g., parentage over or survival of offspring) of intergroup fight *outcomes* are essential to testing theories of the evolution of warfare (4, 5), yet are exceptionally rare outside of human societies (6, 7). In banded mongooses, estrous females benefit from their group engaging in intergroup fights because these females gain fitness benefits in the form of outbred offspring that are healthier than offspring sired by more closely-related within-group males (14, 20). Our analyses suggest that within-group males—who are the main participants in fights and pay most of the costs of competition (13, 14, 18)—are motivated to fight to defend estrous females from mating with males from rival groups (Fig. 2). However, males from groups that won contests in a behavioral sense—repelling their rivals from the site of the fight—could suffer a net loss of fitness (paternity over juveniles) to males from losing groups (Fig. 3). Because we designed our analyses to maximize certainty that any paternity change was a result of a single contest, changes in paternity were rare (12/36 unique contests, 36/197 juveniles), and this result was not statistically significant after controlling for repeated measures of litter, contest, and group identity (Table S5). However, the rate of extra-group paternity we discovered (18% of juveniles sired) is greater that that of most social mammals (mean extra-group paternity rate across 26 species of social mammal = 15.15%; see ref (29)), showing the relevance of intergroup fighting to outgroup mating in this system. More broadly, our finding that gaining extra-group offspring does not come as a result of winning intergroup contests runs counter to theory and data on the evolution of warfare in human societies, where victory in intergroup contests is assumed, or found, to bring fitness rewards in terms of enhanced reproductive success (4, 5, 30). For example, in Nyangatom societies of Ethiopia and South Sudan, victory in intergroup raids results in material and, correspondingly, fitness benefits to male participants (6). By contrast, our finding highlights that contests can result in a disconnect between group success and individual fitness, and that the opportunity costs involved in fighting can undermine group cohesion and performance.

Our results may differ from those of other systems for several reasons, suggesting broader work toward understanding links between conflict outcomes and fitness outcomes. For example, in systems like the Nyangatom and other human (7) and non-human animal systems (e.g., chimpanzees (31)), the same individuals both initiate and participate in conflicts. By contrast, in banded mongooses, females are “exploitative leaders”, initiating contests while males do most of the fighting, including paying most of the costs (14). Similar conflicts of interest in contest participation also occur in other non-human animal systems (32). In vervet monkeys, for example, females motivated to gain food attempt to coerce their male group-mates to fight (33), while males—especially those who are wounded and unable to fight well—punish these coercive females to decrease the chance a fight occurs (34). Perhaps systems with disconnects between conflict initiation and participation are more likely to also have disconnects between conflict and fitness outcomes, but more data is needed to test this hypothesis.

Further data comparing short-term and long-term fitness consequences of intergroup fights, including in banded mongooses, would also be useful to understanding our results. Our analyses focused on the immediate fitness consequences of contest outcomes; that is, paternity over offspring sired at the time of conflict. However, fitness benefits may not be realized until long after a conflict. In the Waorani of Ecuador, for example, men that participate in intergroup raids together are more likely to marry each other’s kin following cultural ideals (cross-cousin marriages), as compared to men who were not raid partners (7). However, these marriages occurred an average of 7 years after the partners’ first raid (7). Similarly, the link between conflict outcomes and fitness outcomes in Nyangatom warriors (see above) is only realized later in life, not at the time of conflict (6). One way that banded mongooses could experience delayed fitness benefits is if certain males participate in conflicts—that is, contribute to collective fighting instead of attempting outgroup mating—in order to signal to within-group females their quality as mates. These males may then receive long-term fitness benefits (i.e., later ingroup matings) that we were unable to quantify here. However, while there is evidence that close inbreeding is avoided within banded mongoose groups (19), our behavioral observations suggest that females show no choice over their male mate guards, and we have no evidence that females select within-group sires based on prior contest participation or fighting ability. In general, models of the evolution of intergroup conflict have not explored variation in the timing or durability of the fitness impacts of victory or loss (4, 5, 30), and understanding variation in short– and long-term fitness consequences could be useful to understanding how selection acts on conflict participation in group-living animals.

Our findings highlight the collective action problem (CAP) that is inherent in intergroup conflict and that occurs more generally in both human and non-human groups (35–37). Collective action problems arise when individuals contribute to a public good (such as winning an intergroup contest) at personal cost, leading to a temptation to defect or shirk. In heterogeneous groups, weaker group members are predicted to free-ride on the contributions of their stronger group-mates, gaining the benefits of group success while minimizing their own efforts (22, 35). Collective action is further undermined if free-riders can not only avoid the costs of contribution, but also gain a fitness benefit by not contributing, as we found is the case for young adult male banded mongooses (Fig. 4). Older males may make up for their free-riding group-mates by contributing more to the fight; indeed, our results (Fig. S1, Table S2) support those of a prior analysis (13) showing that the oldest males in groups have a disproportionate impact on contest success. A recent theoretical model (38) suggests that stronger (here, older) males may “solve” the CAP by overcompensating for lack of investment by weaker individuals, ensuring that heterogeneous groups can remain competitive despite free-riders. Our results are consistent with the predictions of this model: when fighting against groups with estrous females, groups without estrous females (i.e., those facing a potential CAP) still won 38% of contests (Fig. 2).

Much of the recent interest in intergroup conflict stems from its proposed role as a force favoring the evolution of cooperation in group structured populations. For example, the parochial altruism model suggests that the potential for intergroup conflict can lead to the coevolution of intense hostility (parochialism) towards outgroups and intense favoritism (altruism) towards members of an ingroup (4). Banded mongooses have been cited as consistent with this model because of their group structuring (10, 19, 39), which is expected to lead to both high levels of hostility and high levels of within-group cooperation. However, the parochial altruism model (and other evolutionary models of human warfare (5, 30, 40)) assumes that groups contain *n* identical individuals, such that groups do not face internal conflict over whether to initiate contests nor collective action problems over contributions to fighting. The banded mongoose system suggests that within-group heterogeneity can have important effects on the frequency and effectiveness of fighting. The presence of exploitative leaders (females) who take a disproportionate share of the fitness benefits of victory and/or pay disproportionately few costs is predicted to increase the frequency and severity of conflict (14, 41). At the same time, our results show that heterogeneity in the costs and benefits of participation—especially the potential gain of fitness by avoiding fighting and instead mating—can also undermine the effectiveness of collective fighting, making victory less likely. Future theory development and testing on tractable systems would help to reveal how these different factors interact to influence intergroup fighting and the evolution of altruism.

In summary, our results show how fitness benefits to individual group members can have the simultaneous effects of motivating participation in intergroup fights for one group while detracting from participation in the other group. A greater focus on the individual fitness benefits of intergroup contest outcomes will be useful in efforts to test key theories of the evolution of intergroup conflict.

## Methods

### Ethical Note

All research procedures received prior approval from the Uganda Wildlife Authority, the Uganda National Council for Science and Technology, and the Ethical Review Committee of the University of Exeter. All procedures adhere to the ASAB/ABS Guidelines for Animals in Behavioral Research and Teaching.

### Study Population and Data Collection

We collected data on naturally-occurring contests between banded mongoose groups on and around the Mweya peninsula in Uganda (0°12’S, 29°54’E) from February 2000 until April 2019. Following (10), we defined an intergroup contest as occurring when two groups sighted each other and responded by emitting collective “war cry” calls, chasing, and/or engaging in physical fighting. We only analyzed contests in which a clear winner and loser could be determined—losing groups were those that left the area of the contest, while winning groups remained (often chasing away the losing group). Additionally, because we were interested in how male group members (who disproportionately affect contest success (13)) value resources, we only analyzed contests in which each group had at least one male member. Groups without males are very rare and are almost always short-lived; our filtering removed 51 contests involving 3 all-female groups, most (48/51) involved a single all-female group.

For each contest, daily observations of group members allowed us to quantify female estrous status based mainly on observations of mate-guarding: if males were closely mate-guarding females on the day of the contest, the group was considered in group estrus. However, estrus can be subtle and not involve conspicuous mate guarding; therefore, we added data on group estrus by back-calculating from birth data. If any females in a group gave birth within 55-70 days of an intergroup contest, we considered the group to be in group estrus on the date of the contest (55-70 days is a likely timeframe between group estrus and when juveniles are born (15)). The main dataset contained full data on 261 contests between 16 focal groups and 15 rival groups. We also conducted an analysis of the effect of territory location on contest outcomes, using a smaller dataset (given limitations of location data). This analysis is reported in the Supplementary Information; in short, contest location did not affect outcomes.

### Statistical Analysis

All analyses were completed in R software version 4.2.1 (42).

We first identified 104/261 contests where one group was in estrous and the other group was not, treating the estrous group as the “focal” group. In situations where both groups were in estrous (15/261 contests) or neither group was (142/261 contests), we randomly selected a focal group. This random selection was based either on the group the field team was following at the time of the contest (followed group being the focal) or by randomly choosing one of the two competing groups to be the focal using the “sample” function in R. We built a statistical model predicting the probability that the focal (estrous) group won from relative estrous status (both no; one yes, one no; both yes), along with random effects of focal and rival group identity. We fit the model as a Bradley-Terry model (43) in the MCMCglmm package (44). The Bradley-Terry model structure accounts for the fact that, in analyses of the effect of relative (i.e., focal-rival) predictors on contest outcomes when there are repeated observations for each group, focal and rival group identity can be arbitrary and only one observation (win or loss) is recorded for both groups. In these scenarios, a group that consistently wins contests would be expected to win when acting as the (arbitrarily designated) focal group, and lead to losses in a focal group when acting as the (arbitrarily designated) rival group. Therefore, in these models the random effect variance of focal and rival group identity should be equal, and their correlation equal to –1 (see (45–48) for more justification).

Following a similar approach as (48), each model had uninformative parameter-expanded priors and ran for 1,050,000 iterations, where the first 50,000 iterations were discarded and samples were saved every 250 iterations. We ensured good model fit by inspecting plots of effect sizes across iterations, checking that autocorrelation was low among consecutive thinned observations and fixed effects, and by ensuring that Heidelberg and Geweke diagnostic values and plots met expectations for good model fit.

To analyze changes in paternity according to contest outcomes (Fig. 3, Tables S4, S5), we used previously-constructed paternity data for our population (parentage assignments following methods in (49, 50); pedigree deposited as part of data) to identify 1720 juveniles in 388 litters sired by 250 males. We identified the group to which each juvenile was born and the group to which each juvenile’s sire belonged at the time of the juvenile’s birth. We then randomly assigned focal and rival group identity without considering relative estrous status; that is, in scenarios where one group was in estrous and the other was not, we did not assign focal status to the estrous group but instead randomly selected one group as focal. We then identified juveniles born to either the focal or rival group and born within 55-70 days after a contest (a timeframe in which juveniles could likely be sired by members of the rival group (15)). We subset these data to only juveniles born when there was a single contest recorded within the 55-70 day estrus window, and for which we knew which group won that contest. Although we do not observe all contests that groups engage in, this approach gave the most certainty that any change in paternity was a result of a single contest of interest. For each contest, we noted the estrous status of each group (both groups in estrous or only one group in estrous), which group won, and the number of juveniles born to and sired by either competing group. When then built a statistical model (LMM) testing whether focal groups that won intergroup contests gained or lost paternity over a greater number of juveniles. We also included an interaction between focal group outcome (win/loss) and relative estrous status (both groups or only one group in estrous), testing whether the paternity benefits (or costs) of winning changed according to the estrous status of each group. Finally, we included random effects of (1) the contest, (2) the group in which the juvenile’s sire resided at the time of the contest, and (3) the group into which the juveniles were born. By investigating a histogram of model residuals and using the outlierTest function in the car package (51), we identified one outlier in the dataset—a contest in which the focal group won, yet lost paternity of 9 juveniles to males of the rival group. However, because our observations noted no irregular behaviors of group members during the contest, we deemed it a biologically relevant observation and retained this observation in the model. We tested for significant effects of focal group outcome, relative estrous status, and their interaction using a chi-squared test in the Anova function in the car package.

We analyzed the relationship between male age rank and siring success (Fig. 4, Table S7) by first identifying how many juveniles in a given litter were sired by each male. We also calculated each male’s age rank in their own group: we ranked each male in each group according to its age 70 days before a given juvenile was born (i.e., on the earliest date of its conception), with the oldest males in a group holding rank one and males of the same age having the same rank (i.e., ties were allowed). We also categorized whether each juvenile born was sired by an ingroup male—a male residing in the same pack as the juvenile 70 days before the juvenile’s birth—or an outgroup male—a male that was not in the juvenile’s pack 70 days before the juvenile’s birth. We have never observed outgroup mating to take place outside of the context of intergroup contests (*personal observation*); therefore, we assumed that all juveniles born to outgroup males were conceived during intergroup conflict. In the lme4 package (52), we built a generalized linear mixed model (GLMM) with a binomial error structure predicting the proportion of pups sired in each litter from the age rank of male sires, whether the sire was an ingroup or outgroup male, and the interaction of these terms. We also included random effects of sire group identity, sire identity, and litter identity. We ensured good model fit by investigating a histogram of model residuals. We tested for the significance of the interaction between sire age rank and ingroup|outgroup status through a chi-squared test using the Anova function in the car package.

## Authorship statement

Conceptualization, P.A.G., F.M., H.J.N., M.A.C., F.J.T.; Methodology, P.A.G., D.W.E.S., T.C., F.J.T.; Formal Analysis, P.A.G.; Data Curation, P.A.G., D.W.E.S., H.J.N., F.M., F.J.T.; Writing—Original Draft, P.A.G.; Writing—Review & Editing, P.A.G., H.J.N., M.A.C., F.J.T.; Visualization—P.A.G., M.A.C.; Supervision—F.M., M.A.C.; Project Administration—F.M., M.A.C.; Funding Acquisition—P.A.G., M.A.C.

## Data Accessibility Statement

The data and code used in this study have been deposited at Figshare: https://figshare.com/s/53ccc89c078a245e3b69. DOI will be made available as of the date of publication.

## Competing Interest Statement

The authors declare no competing interests.

## Acknowledgments

P.A.G. was funded by Human Frontier Science Program Fellowship LT000460/2019-L and by UC Santa Barbara. The long-term project was supported by National Environment Research Council Grant NE/S000046/1. F.J.T. was funded by a NERC Independent Research Fellowship NE/V014471/1. We thank the Uganda Wildlife Authority and the Uganda National Council for Science and Technology for permission to carry out our research. We thank the Wardens of Queen Elizabeth National Park for logistical support. Solomon Kyabulima, Kenneth Mwesige, Robert Businge, and Solomon Ahabyona helped collect data in the field. We are grateful to Harry Marshall and Emma Vitikainen for curation and maintenance of the long-term data, and Jason Gilchrist, Sarah Hodge, Matthew Bell, Corsin Müller, Neil Jordan, Bonnie Metherell, Roman Furrer, David Jansen, Jenni Sanderson, and Beth Preston for valuable contributions to the project. Members of the University of Exeter Centre for Ecology and Conservation contributed helpful feedback. Jarrod Hadfield, Tom Houslay, Erik Postma, and Alastair Wilson helped with the statistical analyses.

## Supporting Information

### Supplementary analysis—effect of group estrus in context

Our results (Fig. 2) suggest that group estrous impacts conflict outcomes. We further probed this effect by comparing it to the effects of previously-established metrics of RHP and other potential metrics of resource value in this system. The previously-established metrics of RHP were the relative (focal – rival) number of males in each group and the relative age of the oldest male in each group, two factors shown to predict contest outcomes in a prior study (1). The other potential metrics of resource value included the relative number of females and relative number of juveniles in each group. Through data collected *via* daily observations (see Methods), we calculated the number of males, females, and juveniles (group members < 6 months old) in each group on the day of the contest. We also calculated the age of the oldest male in each group as the number of days between the date of the contest and the date of birth of the oldest male in each group. We then selected a focal and rival group for each contest; these focal and rival designations were the same as those described in the Methods for the analysis presented in Fig. 3.

We built a statistical model in which the dependent variable was whether the (randomly-selected) focal group won or lost the contest (binary: 1 = focal win, 0 = focal loss) and the predictor variables were relative proxies of RHP (relative number of males, relative oldest male age) and RV (relative number of females, relative group estrous status, and relative number of juveniles). This model structure—especially the use of randomly selected focal and rival groups—let us as closely as possible compare the effect of group estrous to those of our previously-established RHP metrics as published in (1).

In most cases, variables were integer variables calculated as focal group value *minus* rival group value (e.g., if the focal group had 4 females and the rival group had 2 females, relative N females = 2). The group estrous status variable was similar to that of the model presented in the Main Text; however, instead of testing from the perspective of the group in group estrous (as in the Main Text model), we tested from the perspective of the randomly-selected focal and rival groups. As a result, relative group estrous was a four-level categorical variable with all combinations of focal and rival group estrus: focal group in group estrus and rival group not, focal group not in group estrus and rival group in, both not in group estrus, both in group estrus. Correlations between integer predictor variables were low (Pearson correlation mean = 0.39, range = 0.31, 0.61).

Our statistical model had the same Bradley-Terry structure as that presented in the Main Text, to account for the random selection of focal and rival groups. We also used the same checks of model fit as presented in the Main Text.

In this model, group estrous status was the only resource value metric that strongly impacted contest outcomes (Fig. S1). Focal groups had higher contest success rates when they were in group estrus and their rivals were not. The mean estimate value of this effect was greater than one previously-described RHP metric, oldest male age (Fig. S1; Table S2). However, the relative estrous estimate had a large credible interval. This large CI could be a result of either 1) a truly variable effect of relative estrous on contest success, or 2) a relatively small sample size of contests in which this relative estrous status occurred (56/261 total contests). We used a sensitivity analysis of the RHP effects of relative number of males and relative oldest male age to test whether sample size effects drove this high CI.

We first used our full dataset of 261 contests to build a MCMCglmm model predicting the likelihood of a focal win from the two RHP predictors: the relative number of males and relative age of the oldest male in the contest. This “full dataset model” established the estimate and 95% CI value of both previously-established RHP predictors on the full dataset. Next, we randomly sampled (without replacement) 56 contests from the overall (N = 261 contest) dataset and built a MCMCglmm model of the same structure used on the full dataset. This “downsampled dataset model” established an estimate and 95% CI for both these predictors on a dataset equal in sample size to that for the number of contests in which the focal Y | rival N estrous category occurred. We extracted the mean and 95% CI estimate values from the downsampled dataset model, then repeated this permutation procedure for a total of 1,000 estimate values. We removed models with poor fit, as evidenced by visual inspection of representative “caterpillar plots” and using diagnostics described in the Main Text. Our final datasets included 902 estimates for relative number of males and 915 estimates for relative age of the oldest male.

We plotted these estimates for both the full dataset model and each iteration of the downsampled dataset model (Fig. S1). In total, 63 of 902 (7.0%) of the 95% CIs of the relative number of males effect overlapped zero, while 584 of 915 (63.8%) of the 95% CIs of the relative age of the oldest male effect overlapped zero. Because these previously-established (1), strong predictors of RHP also showed high levels of variability, including 95% CI overlap with zero, in many iterations, it suggests that the large CI of the effect of focal Y | rival N estrous on contest outcomes is a product of a relatively small sample size of contests in which this relative estrous status occurred, not that the effect of relative estrous status is inherently more variable than that of other effects.

### Supplementary analysis—location effects

In many animal societies, the relative location of the contest on each group’s territory impacts contest success (2–4). While we aimed to test this hypothesis alongside the others listed above, location data was only available for 99 contests sampled across 12 years, compared to the 261 contests and nearly 20 years of sampling for the full dataset. When using this limited location dataset and building a model with the full set of predictor variables (i.e., those in Fig. S1), we were unable to achieve good model fit. Therefore, in addition to the statistical model described in the Main Text, we also built a model analyzing the effect of our previously-described metrics of RHP (1) (relative number of males and relative age of the oldest male) along with the relative location of the contest on each group’s territory. This analysis explicitly tests the effect of contest location on outcomes, in as similar a way as possible to our analysis above, but while accounting for dataset limitations.

Our dataset with contest location data (hereafter, the “location” dataset) was a subset of the main dataset that included 99 contests between 8 focal and 9 rival groups. Location data consisted of GPS location fixes of each group, collected by two methods. Between 2006 and 2014, location data were collected at the start and end of every group observation session using handheld Psion II data loggers (model LZ) and GPS units (Garmin eTrex). These data were called “hand” data. Between 2014 and 2019, GPS fixes were collected every thirty minutes (between 0700 and 1900hrs, with a break between 1200 and 1500hrs, corresponding to when groups rest during the heat of the day) using GPS collars (Gipsy4 and Gipsy5, Technosmart, Italy) fitted to a maximum of two individuals in each group. These data were called “collar” data. The GPS collar data were filtered for accuracy following methods in(5). In addition to collecting location data throughout each day, the location of each contest was collected (either by hand or collar) at the time of the contest.

Our filtering process removed several GPS locations prior to analysis. GPS locations were discarded if they fell below four satellite connections or if their “horizontal dilution of precision” values (accounting for heteroscedasticity from satellite reception) exceeded a value above four (6). GPS locations were also discounted if they exceeded unrealistic altitudes (<800m or > 1100m) (7). All GPS data were clipped using a detailed map of the Mweya peninsula’s shape, to discount locations that fell in nearby Lake Edward, which is impassable for banded mongooses. GPS collar data often contained groups with two individuals collecting GPS data at one time. Including these records would constitute a duplicated timestamp and bias home ranges (8). Instead, we kept only records for the mongoose which registered the greatest number of GPS locations. We did not average, or try to keep both of, the duplicated GPS trajectories because other days had only one collar measuring group location, and therefore this method is consistent with these non-duplicated days. Finally, we removed GPS locations where individuals wearing collars were guarding juveniles at the den, as any such data would not be reflecting group movement for that time period. In banded mongooses, groups only split when some individuals are left at the den with juveniles (9, 10). Whether individuals left at the den were also wearing a GPS collar was assessed via field observations.

Our measure of the location of a contest on each group’s territory involved calculating how much time each group spent at the contest location before the contest occurred. Using default parameters in the ctmm package (11) we first built utilization distributions (UDs) for the GPS location data of each group involved in the contest for the 90 days preceding the date of the contest (a timeframe representing a full reproductive period (12)). These UDs, which accurately account for autocorrelation in the telemetry data, represent a “familiarity index” of each grid cell (average grid cell size was 5 x 5m). The familiarity index describes how often the group was found in each grid cell, given the observed 90-day locations data. From these UDs, we then used the “raster” function in ctmm to generate a rasterized cumulative distribution function (CDF) of the location data. For each group (focal and rival), we calculated the rasterized CDF value of the location of the contest and subtracted this CDF value from one. This created a metric in which a value of 1 represented a grid of the group’s UD that had been the group’s most occupied grid cell for the 90 days preceding the contest, while a value of 0 represented an unoccupied area of the group’s UD in the same timeframe. We subtracted the rival group’s value from the focal group’s value to create a relativized metric that we called “relative familiarity”. If relative familiarity for a given contest was greater than 0, it meant the focal group spent more time at the location of the contest in the preceding 90 days than the rival group. Alternatively, if relative familiarity was less than 0, it meant the rival group spent more time at the contest location in the preceding 90 days as compared to the focal group.

With the location dataset, we built a statistical model predicting the probability that the focal group won the contest from the relative (focal – rival) number of males, relative age of oldest male (in days), and relative familiarity. We built this model in the MCMCglmm package following the same methods, including the same checks of model fit, as those described for the model presented in Fig. S1.

The analysis of location data showed that relative familiarity did not influence contest outcomes (Table S3). Additionally, compared to the datasets used in (1) (their Table S4) and in our analysis of the main dataset (Fig. S1 and Table S2), this model had lower estimates for the effects of both number of males and oldest male age on contest outcomes (Table S3). We found no differences in the distributions of raw data on number of males or oldest male age among the datasets (dataset used in Main Text, location dataset, and dataset from (1); pairwise t-tests comparing number of males data among datasets, all P > 0.30; comparing oldest male age data, all P > 0.18). Instead, this discrepancy may be a result of decreased date ranges sampled in the location dataset. The dataset used in the Main Text sampled 19.7 years (7205 days), the dataset used in (1) sampled 15.4 years (5622 days), and the location dataset sampled only 12.3 years (4493 days).

Overall, we found no evidence that the location of the contest on each group’s territory influenced contest outcomes.

### Supplementary analysis—effect of contest outcomes on proportion of litter sired

We asked whether the finding that winning groups tended to be more likely to lose paternity over juveniles, as compared to losing groups (Fig. 3), also held when we accounted for the total number of juveniles born in each litter. Following similar approaches as the model presented in Fig. 3, we built a binomial GLMM in which the outcome variable was the proportion of juveniles in a given litter of which paternity was gained, or retained, by the focal group. That is, “successes” in this binomial model were juveniles over which paternity was gained or retained, while “failures” were juveniles of which paternity was lost to males in the rival group. The predictor variables included focal outcome (win/loss) and relative estrous status (one yes | one no, both yes), and random effects included contest and group identity (for both the group siring the juveniles and the group into which the juveniles were born). Note that, in contrast to the model presented in the Main Text, we did not include an interaction term between focal outcome and relative estrous status. We removed this interaction term after identifying high variance inflation factors in predictor variables when the interaction was included (13).

This proportional model found the same directionality, and similar strength, of effects as the model presented in the Main Text (Table S6). Focal groups that won contests sired a lower proportion of juveniles per litter, as compared to focal groups that lost contests. Therefore, the finding that behavioral wins are not positively correlated with fitness gains holds not only for the overall number of offspring sired (Fig. 3), but also when considering the overall number of offspring available.

## Supporting Information Figures

**Fig. S1.**
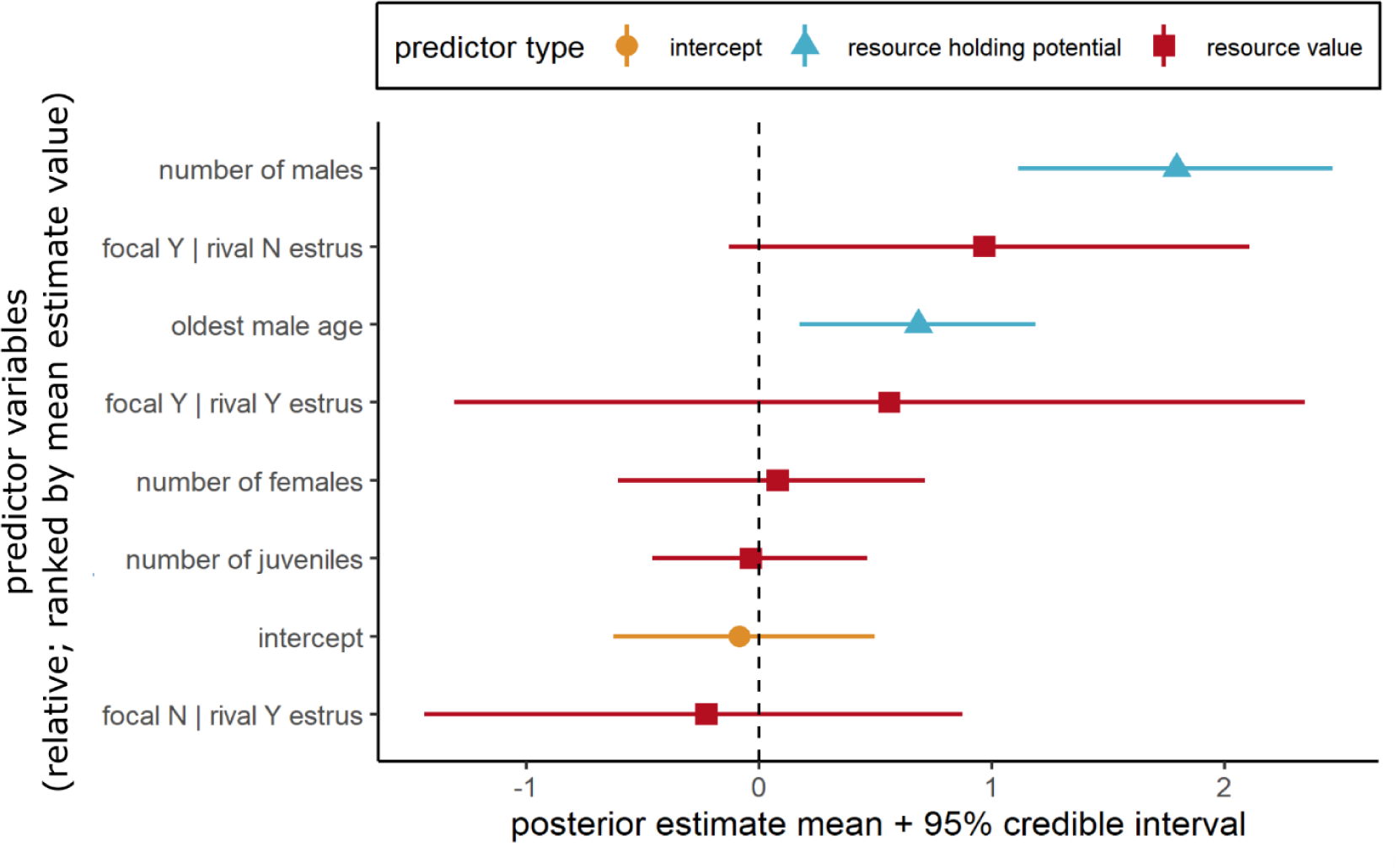
When focal groups were in group estrus and rival groups were not (focal Y | rival N estrus), focal groups had higher contest success. The value of this estimate was second-highest in the model, though small sample size led to a high credible interval (Table S1, Fig. S2). Plot shows posterior estimates from MCMCglmm model testing how resource holding potential and resource value metrics affected contest outcomes. On x-axis, points represent estimate means, lines represent 95% credible intervals. Predictor variables (y-axis) are arranged in decreasing order of estimate mean.

**Fig. S2.**
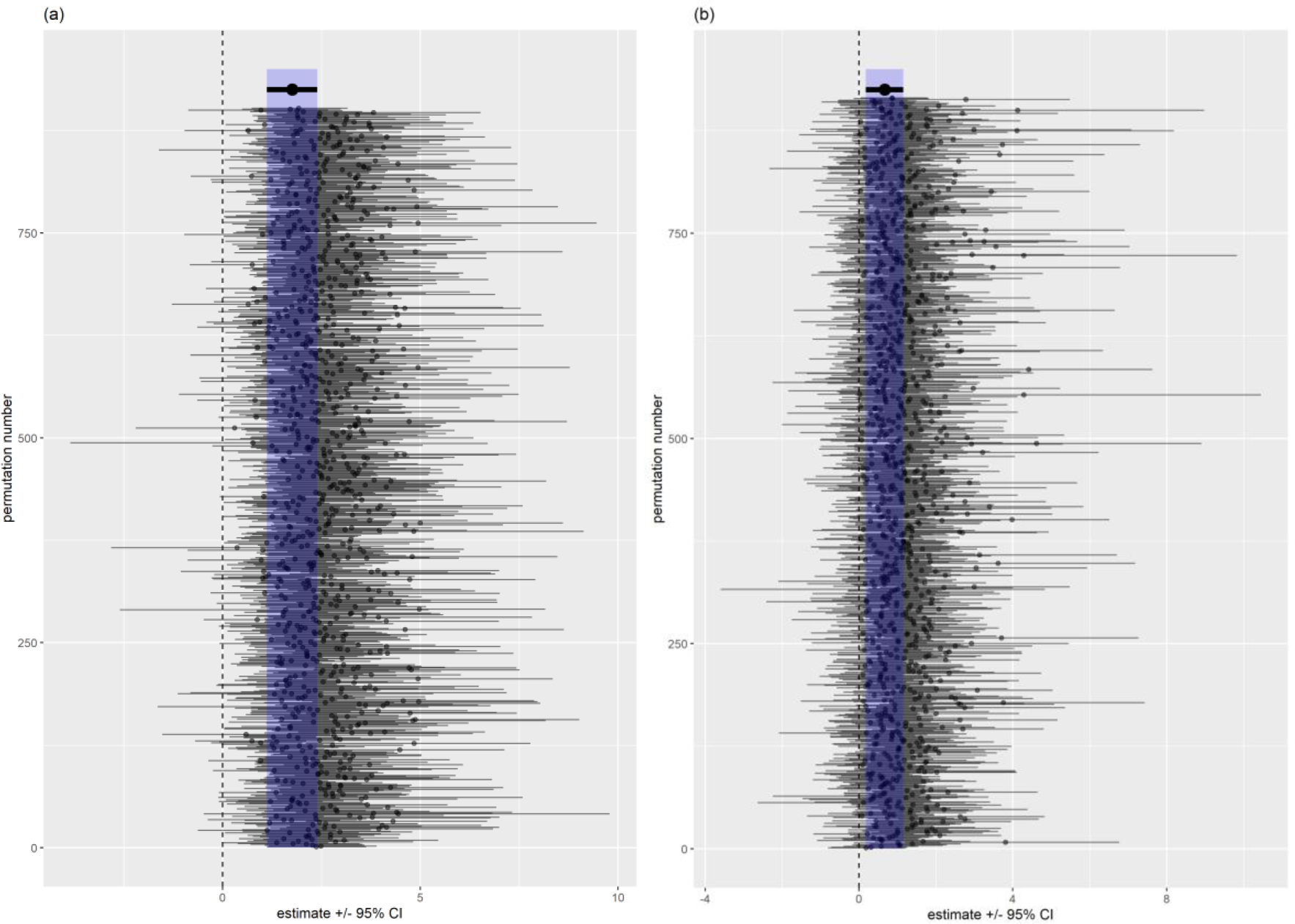
Plot showing estimates and 95% credible intervals for permutation analyses of the effect of (a) relative number of males and (b) relative age of the oldest male on contest success. At top of each figure, the estimate (point) and 95% CI (lines) for the effect of relative number of males and oldest male age is shown from the full dataset (N = 261). Blue shaded region shows the 95% CI of this full-dataset analysis throughout the plot. Small points and bars show estimate and 95% CI (x-axis) for each of ∼1,000 MCMCglmm models predicting contest success from both relative number of males and relative oldest male age, in which the sample size of contests was made equal to that of the focal Y | rival N relative estrous category in the model presented in Fig. S1 (56 contests). Dashed vertical line shows estimate value of 0; small bars crossing this line represent models in which the pMCMC of each effect would be greater than 0.05.

## Supplementary Tables

**Table S1.** Output from MCMCglmm model predicting estrus group win/loss from relative estrus status. Estimates for relative estrus categories are in comparison to the “Both no” category. Asterisks in pMCMC column show p < 0.05.

| Predictor | Posterior estimate mean | 95% CI of posterior estimate | pMCMC |
| --- | --- | --- | --- |
| Intercept | -0.11 | -0.50, 0.30 | 0.60 |
| One yes, one no | 0.69 | 0.08, 1.33 | 0.03* |
| Both yes | 0.60 | -0.64, 1.92 | 0.36 |
| Random effect | Posterior mean | 95% CI of posterior mean |  |
| Estrus focal ID + estrus rival ID | 0.08 | 0.00, 0.34 |  |

**Table S2.**
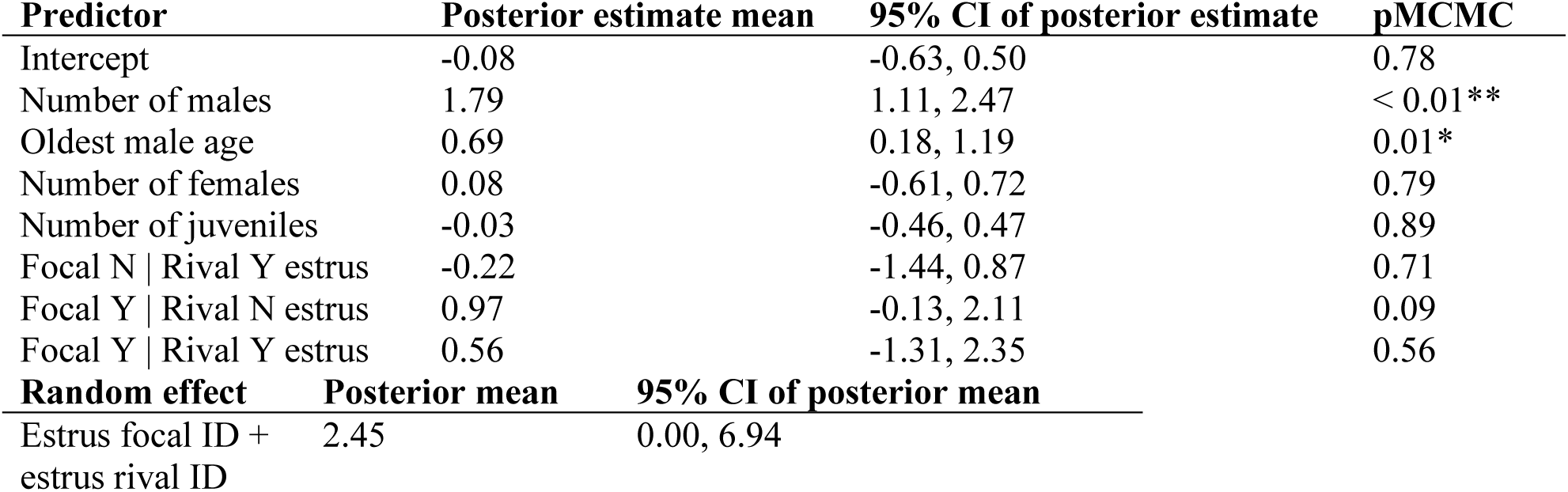
Output from MCMCglmm model predicting focal win/loss from RHP and RV predictors. Estimates for relative estrus categories are in comparison to the “Focal N | Rival N” estrus category. Asterisks in pMCMC column show ** p < 0.01, * p < 0.05.

**Table S3.** Output of MCMCglmm model predicting contest outcomes from RHP variables and relative familiarity (representing contest location). Asterisk shows *pMCMC < 0.05.

| Predictor | Posterior mode | 95% CI | pMCMC |
| --- | --- | --- | --- |
| Number of males | 0.88 | 0.76, 1.84 | 0.046* |
| Oldest male age | 0.35 | -0.39, 1.12 | 0.335 |
| Relative familiarity | -0.12 | -1.05, 0.57 | 0.541 |

**Table S4.** Summary statistics for paternity of juveniles as a result of focal group outcomes and relative estrous category. For each win/loss and estrous category, table lists the number of juveniles of which paternity was gained or lost by the focal group, the net difference (gain – loss), and the number of juveniles of which paternity was retained. Other columns list the number (N) of unique contests, litters, and focal and rival groups for each category.

| Focal loss or win | Estrous category | Paternity gained | Paternity lost | Net paternity (gain – loss) | Paternity retained | N unique contests | N unique litters | N unique focal groups | N unique rival groups |
| --- | --- | --- | --- | --- | --- | --- | --- | --- | --- |
| Loss | Both yes | 4 | 0 | 4 | 4 | 4 | 5 | 4 | 3 |
|  | One yes, one no | 2 | 6 | -4 | 14 | 15 | 15 | 9 | 4 |
| Win | Both yes | 1 | 6 | -5 | 4 | 3 | 5 | 3 | 2 |
|  | One yes, one no | 5 | 12 | -7 | 16 | 17 | 17 | 6 | 8 |

**Table S5.** Output from model predicting the gain or loss of paternity by focal groups from contest outcome, estrous status, and their interaction.

| Fixed effect | Contrast | $\beta$ | SE | $\chi^2$ (df) | p |
| --- | --- | --- | --- | --- | --- |
| Intercept |  | 0.84 | 0.91 | 0.86 (1) | 0.35 |
| Focal outcome | Loss | 0.00 | 0.00 |  |  |
|  | Win | -1.23 | 0.80 | 2.36 (1) | 0.12 |
| Estrous category | Both yes | 0.00 | 0.00 |  |  |
|  | One yes, one no | -0.65 | 0.67 | 0.93 (1) | 0.33 |
| Focal outcome : estrous category | Both yes | 0.00 | 0.00 |  |  |
|  | One yes, one no | 1.02 | 0.94 | 1.18 (1) | 0.28 |
| Random effect | Variance | SD |  |  |  |
| Contest ID | 0.00 | 0.00 |  |  |  |
| Birth group ID | 1.72 | 1.31 |  |  |  |
| Sire group ID | 2.09 | 1.45 |  |  |  |

**Table S6.** Output from model predicting the proportion of juveniles per litter over which paternity was retained or gained by the focal group from contest outcome and estrous status.

| Fixed effect | Contrast | $\beta$ | SE | $\chi^2$ (df) | p |
| --- | --- | --- | --- | --- | --- |
| Intercept |  | 11.59 | 6.69 | 3.00 (1) | 0.08 |
| Focal outcome | Loss | 0.00 | 0.00 |  |  |
|  | Win | -4.19 | 2.57 | 2.66 (1) | 0.10 |
| Estrous category | Both yes | 0.00 | 0.00 |  |  |
|  | One yes, one no | 3.20 | 3.50 | 0.84 (1) | 0.36 |
| Random effect | Variance | SD |  |  |  |
| Contest ID | 1.65 | 1.29 |  |  |  |
| Birth group ID | 39.57 | 6.29 |  |  |  |
| Sire group ID | 38.54 | 6.21 |  |  |  |

**Table S7.** Output from model predicting the proportion of juveniles in each litter from sire age rank, ingroup|outgroup status, and their interaction. Asterisks show *p < 0.05 and **p < 0.01

| Fixed effect | | $\beta$ | SE | $\chi^2$ (df) | p |
| --- | --- | --- | --- | --- | --- |
| Intercept |  | -0.88 | 0.06 |  |  |
| Sire age rank |  | -0.05 | 0.01 | 17.49 (1) | < 0.01** |
| Ingroup \| outgroup | Ingroup | 0.00 | 0.00 | 8.15 (1) | < 0.01** |
|  | Outgroup | -0.45 | 0.13 |  |  |
| Sire age rank : ingroup \| outgroup |  | 0.04 | 0.02 | 4.90 (1) | 0.03* |
| Random effect | Variance | SD |  |  |  |
| Sire group | 0.04 | 0.06 |  |  |  |
| Sire ID | 0.01 | 0.12 |  |  |  |
| Litter ID | 0.06 | 0.24 |  |  |  |

## Notes

https://figshare.com/s/53ccc89c078a245e3b69

### Competing Interest Statement

The authors have declared no competing interest.

